# Assessment of 3D MINFLUX data for quantitative structural biology in cells revisited

**DOI:** 10.1101/2022.05.13.491065

**Authors:** Klaus C. Gwosch, Francisco Balzarotti, Jasmin K. Pape, Philipp Hoess, Jan Ellenberg, Jonas Ries, Ulf Matti, Roman Schmidt, Steffen J. Sahl, Stefan W. Hell

## Abstract

Prakash and Curd provide a re-analysis^1^ of individual datasets taken from our report^2^ demonstrating MINFLUX 3D imaging in cells. Their evaluation confirms the unique localization precision provided by MINFLUX^2,3^ featuring a standard deviation of σ = 1-3 nm. We appreciate their confirmation and also welcome the opportunity to clarify their remaining points. The hitherto almost unconceivable 3D localization precision attained by MINFLUX is likely to hold the key to an all-optical dynamical structural biology.

## Introduction

As we have explained in the paper^2^, a fluorescence microscope can image only the distribution of fluorophores in the sample – not the labeled biomolecules per se. Therefore, the concept of resolution can only be applied to the fluorophores. As long as fluorophores are individually identified, the localization precision is tantamount to the resolution. By reaching σ on the size scale of the fluorophores themselves (1-3 nm), our paper shows that such a remarkable imaging performance can indeed be reached in cells.

Microscopy images from many individual nuclear pore complexes (NPCs) inherently reflect the biological variability of the NPC *in cellulo*^4^, including deviations from roundness and 8-fold symmetry^5^. Results gained from averaging over up to a thousand NPCs, as in cryo-electron tomography (ET), cannot represent the ground truth for each and every NPC in the cell. In fact, NPC images are expected to vary in terms of pore diameter and protein positions, because their conformation is affected by nucleocytoplasmic transport state and forces in the membrane^6–8^. The nanometer-level variations in the MINFLUX data of individual NPCs relative to the averaged cryo-ET rendition actually highlight the ability of this nanoscopy method to access the structure of individual protein complexes. In our view this is a strength and not a weakness of the method, as Prakash and Curd imply.

Since fluorescence microscopy renders just the fluorophores, it is obvious that the degree and type of labeling, the biomolecule-fluorophore distance, and the on/off-switching properties have major consequences for what can be biologically inferred. These factors, however, also depend on the mounting medium, fixation, chemical environment and the type of fluorophore employed in the sample^9^. Moreover, as these factors may vary from sample to sample, drawing general conclusions on an imaging method from individual recordings is questionable, as are basically all of the assertions made by Prakash and Curd:

## Results and Discussion

1. *“The average z distance between cyto- and nucleoplasmic layers of Nup96 localizations was 40.5 nm instead of ~50 nm, in the dataset on which this claim was based”*. Unfortunately, in their analysis of our data it appears that Prakash and Curd neglected to account for the local inclination of the nuclear envelope (Figs. 3f,g of Gwosch *et al*.). Calculating a relative position distribution (RPD) along the *z* axis over the entire field of view (see their Fig. 2g-j) cannot yield the correct distance between the two Nup96 rings. It biases the RPD towards lower values. We demonstrate this effect by repeating their analysis (Fig. 1a). If the membrane is assumed to be flat and the nuclear pore tilts are not accounted for – i.e., the 3D dataset of Fig. 3f of Gwosch *et al*. is used as is – the RPD(Δ*z*) indeed peaks at the erroneous value of ~40 nm (Fig. 1a, dashed line) derived by Prakash and Curd. Taking the inclinations of individual NPCs into account shifts the RPD peak to a distance of ~47 nm, close to the expected ~50 nm. Histograms of localizations from the aligned NPCs along the *z* direction (Fig. 1b) exhibit ~50 nm distance between the two layers, as do particle-averaged^10^ data from a separate newly acquired recording of a larger number of NPCs (Fig. 1c).
2. *“The mean or best-fit Nup96 ring diameter varies between datasets and the spread of diameters in each dataset is broader than that found by [PALM/STORM].”* Even if we consider that fixation may add to the variability of the many individual snapshots of NPCs in various samples, the average diameters (107, 108, 111 nm) determined by Prakash and Curd still fall within the respective error margins, and are fully compatible with the 107 nm found by cryo-electron tomography after extensive averaging^11^. Therefore, although they are based on small datasets (*N* = 20 in each case), the diameter measurements actually agree very well.
3. *“The eightfold symmetry of NPCs is rarely visible at a single nuclear pore level and was not clearly determined in structure-based modelling of the localisation datasets*”. We used the formulation to indicate that the individual corners of the well-known 8-fold symmetrical NPC can be identified in the MINFLUX images in general. In fact, due to incomplete labeling a variable number of displayed subunits is expected for each pore. The symmetry is further obscured for many of the 2D images by frequent NPC tilts, as in many other superresolution reports. To account for these tilts, 3D imaging becomes essential. Certainly, camera-based PALM/STORM provides larger fields of view than the scanning-beam MINFLUX demonstrations to date, which is why PALM/STORM can yield many more NPC images in one shot, albeit at lower localization precision. The higher numbers of pores in a field of view facilitate the generation of well-averaged images or the picking of examples that are more completely labeled. Yet we agree with Prakash and Curd that the apparent labeling efficiency in the 3D MINFLUX NPC imaging – not the resolution – was lower than in the referenced STORM study. Among the obvious remedies are labeling optimizations such as replacing the utilized STORM-dye Alexa 647 (which is not optimal for MINFLUX) with MINFLUX-optimized dyes that are currently under development. The lower apparent labeling is more related to the labeling or the fluorophore used than to the MINFLUX concept.
4. *“Furthermore, in 2-color imaging, the inner ring found in similar [PALM/STORM] experiments at 40-nm diameter^12,13^ was not resolved as a ring by MINFLUX”*. In the STORM reports^12,13^ highlighted by Prakash and Curd, rare NPC images were picked by the authors out of many hundreds where a ring is not visible either. Reasons for that may be insufficient labeling, tilts and structural variabilities^7,8^. The visibility of the channel (ring) requires a good top view, which was easier to accomplish in the referenced STORM study^12^ because that study used isolated Xenopus nuclei instead of complete mammalian cells, as we did. Nonetheless, the ring-like structure mapped out by WGA can also be extracted from more recently recorded MINFLUX data (Fig. 1d-g) for selected top (*x*-*y*) views. An overlay of the aligned MINFLUX data showed a narrowly defined WGA ring of ~30 nm diameter in fixed U-2 OS cells. Yet, care should be exerted when taking these ‘rings’ for granted. Considering the overall distribution and accessibility of the putative WGA binding partners, further research will be needed to answer if the observed rings are biologically relevant, or whether they are due to labeling and fixation.
5. *“[Live MINFLUX and filtering.] We reproduced the published images of Gwosch et al. (2020) and noted an increased level of filtering”*. The procedure separating emission events from background was performed without the use of prior information about the imaged structures and following clearly described criteria^2^ based on the so-called central donut fraction, the estimated location of the molecule with respect to the center of the scan pattern, and the photon number in the last MINFLUX iteration. Bearing in mind the pioneering nature of our experiments, much more data was saved at the time (as a precaution) than eventually had to be used. This naturally led to a large fraction of excluded events in post-processing. More recently published MINFLUX imaging introduced convenient elements of real-time event classification already at the stage of data acquisition^14^. Finally we note that the live-cell MINFLUX images of Nup96 fused to the fluorescent protein mMaple (Fig. 2f in Gwosch *et al.),* outperform similar attempts by PALM.

**Fig. 1.**
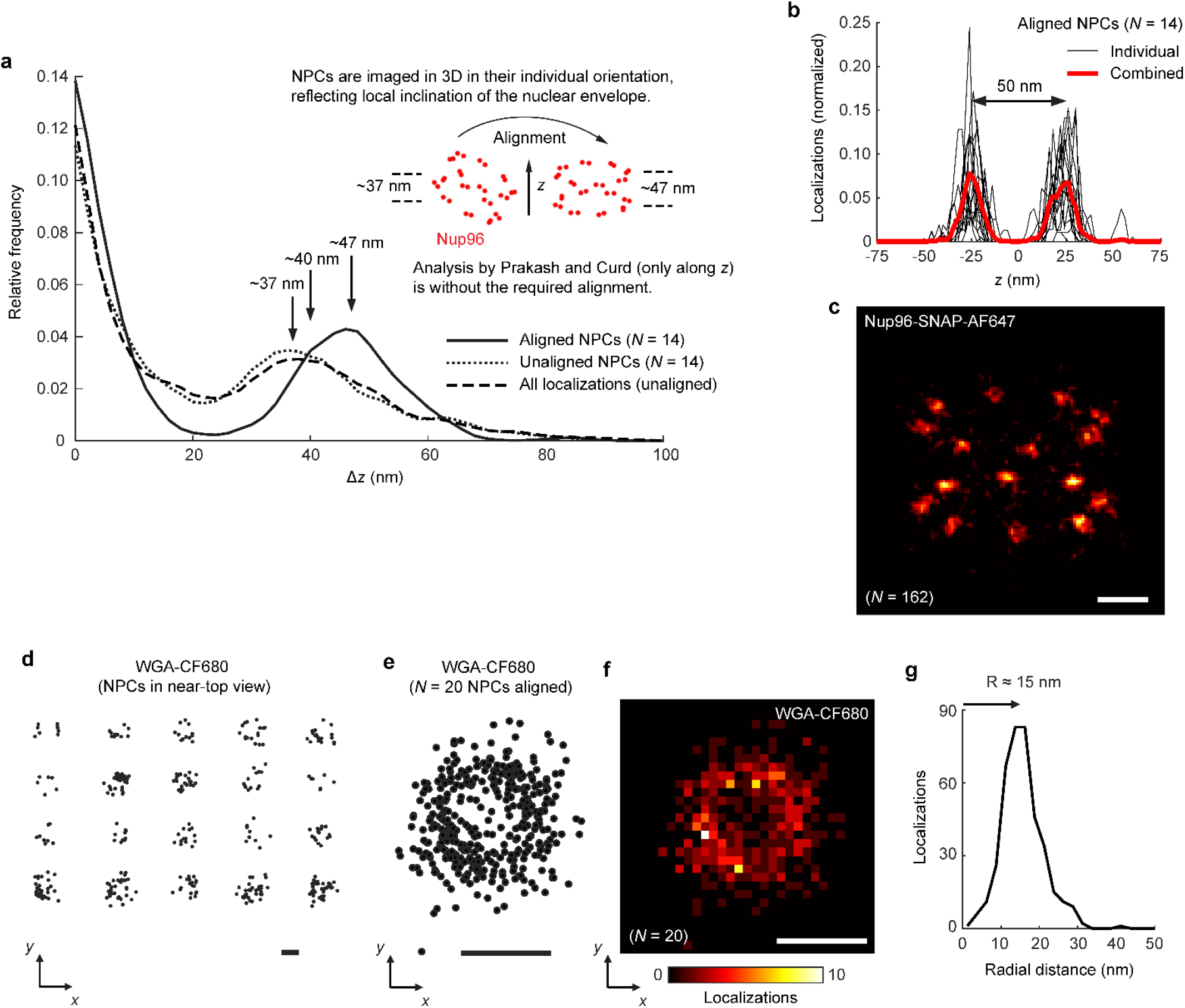
(a-c) Prakash and Curd neglected the tilts of individual nuclear pore complexes in non-flat nuclear membranes. An analysis based on the relative position distribution (RPD) along the *z* axis is inadequate for extraction of the inter-layer distance (ring-to-ring spacing) from Nup96 3D localizations, unless the data from individual NPCs is aligned to fall along the direction of analysis. The individual NPC tilt has to be corrected for in order to enable accurate measurements. (**a**) We estimated the tilts of a subset (*N* = 14) of NPCs for which the labeling was sufficient such that the orientation could be inferred by a 3D fitting analysis. The RPD(Δ*z*) for the unaligned NPCs peaks at ~37 nm (dotted line), reflecting the presence of non-negligible tilts. After their reorientation to the horizontal, the resulting RPD(Δz) of the NPCs peaks at ~47 nm (bold line), which is close to the ~50 nm expected from cryo-electron tomography. (**b**) Histograms of Nup96 localizations from the aligned NPCs along the *z* direction. Black: individual NPCs. Red: 14 NPCs combined. (**c**) Averaging result by particle fusion of Nup96 data from 3D MINFLUX, using the code from ref.^10^ with the prior knowledge of 8-fold pore symmetry. AF: Alexa Fluor 647. Scale bar: 25 nm. **(d-g) MINFLUX data of WGA-CF680 binding patterns exhibit a ring-like structure.** The individual 3D tilts of NPCs obscure the channel in the majority of NPCs. In top views (*x*-*y*), a hollow localization distribution was therefore observed in a small subset of NPCs imaged in chemically fixed U-2 OS cells. (**d**) To illustrate the effect, *N* = 20 NPCs, which exhibited a top view or near-top view, were selected by visual inspection. (**e**) An overlay of the aligned data forms a ring of ~30 nm diameter. (**f**) Histogram of the aligned localizations. (**g**) Radial distribution of localizations, indicating a radius of ~15 nm. Scale bars (d-f): 25 nm.

The choice of fluorophore (and tag) is decisive for the performance of any superresolution method. The utilized dye Alexa Fluor 647 has been *the* go-to STORM dye for one and a half decades. However, due to key aspects of its photophysical behavior, it is certainly not the optimal dye for MINFLUX. Therefore, comparing MINFLUX with STORM without highlighting the limits of Alexa Fluor 647 as a caveat is neither objective nor definitive.

Molecule-specific imaging at molecule-scale 3D resolution is bound to empower biological research. As for any new fluorescence imaging concept, for MINFLUX we expect continuing progress, especially in conjunction with further advances in fluorescence labeling. Multiple MINFLUX setups are operated in leading research laboratories around the world, including EMBL’s new Imaging Centre, which provides open user access to the technology^15^. What’s even more exciting in our view, is the fact that the hitherto almost unconceivable 3D localization precision of 1-3 nm attained by MINFLUX (and confirmed by Prakash and Curd) is likely to hold the key to an all-optical dynamical structural biology.

## Methods

The MINFLUX data shown reanalyzed in Fig. 1a and b was taken from Fig. 3f of Gwosch *et al*.^2^, and the data presented in Fig. 1c and d-g was recorded anew on an Abberior Instruments MINFLUX nanoscope^14^ following previously established fluorescence labeling and imaging protocols^13,14^.

### Cell culture

Before seeding of cells, high-precision 18 mm round glass coverslips (No. 1.5H, Marienfeld, Germany) were cleaned by placing them in ethanol overnight, rinsed with water and dried in a laminar flow cell culture hood before finally irradiating the coverslips with ultraviolet light for 30 min.

Nup96-SNAP cells (300444, CLS, Heidelberg, Germany) were seeded on clean glass coverslips 2 days before fixation to reach a confluency of about 50–70% on the day of fixation. The cells were grown in growth medium (DMEM (11880-02, Gibco)) containing 1× MEM NEAA (11140-035, Gibco), 1× GlutaMAX (35050-038, Gibco) and 10 % (v/v) fetal bovine serum (10270-106, Gibco) at 37°C and 5 % CO_2_. Before further processing, the growth medium was aspirated, and samples were rinsed with PBS (room temperature) to remove dead cells and debris.

### Preparation of NPC samples

Cells on glass coverslips were prefixed in 2.4 % (w/v) FA (formaldehyde) (15710, EMS, Hatfield, PA, USA) in PBS for 20 s before incubating them for 10 min in 0.5 % (v/v) Triton X-100 in PBS. Fixation was completed in 2.4 % (w/v) FA in PBS for 20 min. FA was quenched for 5 min in 100 mM NH4Cl in PBS and then washed 3x for 5 min in PBS. Fixed cells were blocked with Image-IT signal enhancer (Thermo Fisher, Waltham, MA, USA) for 30 min and then incubated with 1 μM BG-AF647, 0.5 % BSA and 1 mM DTT in PBS for 1 h to stain Nup96-SNAP-tag. Cells were washed 3x for 5 min with PBS and subsequently imaged.

For imaging with WGA-CF680 the sample was incubated for 10 min with 1:10.000 diluted WGA-CF680 (29029-1, Biotium, Fremont, CA, USA) in 100 mM Tris, pH 8.0, 40 mM NaCl, and rinsed 3x with PBS shortly before mounting. Samples were mounted on glass slides (12290, bmsmicroscopes.com) with a small cavity to hold the imaging buffer and sealed with 2K silicone (eco-sil, 1300 9100, www.picodent.de).

### Imaging buffer

Glucose oxidase/catalase buffer supplemented with cysteamine (MEA) was used to image Nup96-SNAP-AF647 and WGA-CF680 samples. GLOX+MEA contained 50 mM Tris/HCl pH8, 10 mM NaCl, 10 % (w/v) D-glucose, 500 μg/ml glucose oxidase, 40 μg/ml glucose catalase and 14 mM MEA in H_2_O (all reagents from Sigma).

### Data analysis

The described analysis was performed in MATLAB. In brief, for Fig. 1a the RPD(Δ*z*) was calculated for three separate sets of 3D localization coordinates: (1) All localizations within the field of view, unaligned; (2) 14 segmented NPCs, unaligned; and (3) the same 14 segmented NPCs after horizontal alignment (rigid transformation of their coordinates), removing the tilts estimated from a comparison to a template of the average positions of Nup96 C termini based on the cryo-ET structure^11^. For the localization coordinate set (1), all Δ*z* across the entire field of view were calculated, but adding a filter in the Euclidian distance *R* < 200nm for meaningful interpretation of the RPD histogram, due to the substantial non-flatness of the nuclear membrane over the field of view. Figure 1b shows direct, normalized z histograms of the aligned localizations of set (3). For Fig. 1c, 162 NPCs were segmented from the field of view of a Nup96 recording by visual inspection, and their localization data was averaged using the code of ref.^10^ with the prior knowledge of 8-fold pore symmetry. The combined set of WGA-CF680 *x,y* localizations from 20 NPCs (localizations in Fig. 1e, histogram in Fig. 1f) was obtained by fitting circles to the individual localization patterns of Fig. 1d, followed by co-alignment based on their centers.

## References

1. Prakash, K. & Curd, A.P. Assessment of 3D MINFLUX data for quantitative structural biology in cells. https://doi.org/10.1101/2021.08.10.455294 (2021).

2. Gwosch, K.C. et al. MINFLUX nanoscopy delivers 3D multicolor nanometer resolution in cells. Nat. Methods 17, 217–224 (2020).

3. Balzarotti, F. et al. Nanometer resolution imaging and tracking of fluorescent molecules with minimal photon fluxes. Science 355, 606–612 (2017).

4. Sabinina, V.J. et al. Three-dimensional superresolution fluorescence microscopy maps the variable molecular architecture of the nuclear pore complex. Mol. Biol. Cell 32, 1523–1533 (2021).

5. Löschberger, A., Franke, C., Krohne, G., van de Linde, S. & Sauer, M. Correlative super-resolution fluorescence and electron microscopy of the nuclear pore complex with molecular resolution. J. Cell Sci. 127, 4351–4355 (2014).

6. Pulupa, J., Prior, H., Johnson, D.S. & Simon, S.M. Conformation of the nuclear pore in living cells is modulated by transport state. eLife 9, e60654 (2020).

7. Schuller, A. et al. The cellular environment shapes the nuclear pore complex architecture. Nature 598, 667–671 (2021).

8. Zimmerli, C.E. et al. Nuclear pores dilate and constrict in cellulo. Science 374, eabd9776 (2021).

9. Sahl, S.J., Hell, S.W. & Jakobs, S. Fluorescence nanoscopy in cell biology. Nat. Rev. Mol. Cell Biol. 18, 685–701 (2017).

10. Heydarian, H. et al. 3D particle averaging and detection of macromolecular symmetry in localization microscopy. Nat. Commun. 12, 2847 (2021).

11. von Appen, A. et al. In situ structural analysis of the human nuclear pore complex. Nature 526, 140-143 (2015).

12. Löschberger, A. et al. Super-resolution imaging visualizes the eightfold symmetry of gp210 proteins around the nuclear pore complex and resolves the central channel with nanometer resolution. J. Cell Sci. 125, 570–575 (2012).

13. Thevathasan, J.V. et al. Nuclear pores as versatile reference standards for quantitative superresolution microscopy. Nat. Methods 16, 1045–1053 (2019).

14. Schmidt, R. et al. MINFLUX nanometer-scale 3D imaging and microsecond-range tracking on a common fluorescence microscope. Nat. Commun. 12, 1478 (2021).

15. https://www.embl.org/about/info/imaging-centre/

